# Diversity and Metabolic Potential of Microbial Communities in a Serpentinite-Hosted Spring at the Tablelands

**DOI:** 10.1101/2021.09.15.460147

**Authors:** Emily Dart, William J. Brazelton

## Abstract

The geochemical process of serpentinization releases energy and organic carbon: two of the basic requirements needed to support life. Sites of active serpentinization in the deep subsurface provide the intriguing possibility of a non-photosynthetically-supported biosphere. However, serpentinization also creates conditions, such as high pH and limited electron acceptors, which may limit microbial growth and diversity. Gaining an understanding of the identity and metabolic potential of microbes that thrive in these environments may provide insight as to whether serpentinization is sufficient to independently support life. Tablelands Ophiolite in Gros Morne National Park, Newfoundland, Canada is a continental site of serpentinization where serpentinite springs form surface pools. These pools provide easy sampling access to subsurface fluids and may allow for sampling of the subsurface microbial community. However, identification of members of the subsurface community in these pools is complicated by both surface contamination and contamination by organisms that inhabit the transition zone where hydrogen-rich subsurface fluids meet oxygen-rich surface fluids. This study was designed to distinguish among these potential sources of microorganisms by using a sampling technique that more effectively samples subsurface fluids. Community dissimilarity comparisons using environmental 16S rRNA gene sequencing indicate that the sampling design led to more direct access to subsurface fluids. These results are supported by metagenomic analyses that show metabolic pathways consistent with non-photosynthetic carbon fixation in the samples expected to represent subsurface fluids and that show hydrogen oxidation pathways in samples associated with the surface sources. These results provide a clearer picture of the diversity and metabolic potential of microbial communities potentially inhabiting subsurface, serpentinite-hosted habitats.

## Introduction

Estimates of global biomass indicate that microorganisms may account for up to half of the Earth’s carbon pool, meaning that the amount of carbon stored in microorganisms may be comparable to the amount of carbon stored in plants (Whitman *et al*., 1998). Although many of these microorganisms thrive in the photosynthetically-supported biosphere, the majority of these microorganisms exist in subsurface environments (Whitman *et al*., 1998; Kallmeyer *et al*., 2012). Even though early estimates for the amount of microbial life in some types of marine sediments are now thought to be too high (Kallmeyer *et al*., 2012), the total amount of microbial biomass in all subsurface habitats is still expected to outweigh surface biomass (Edwards *et al*., 2012). However, confidence in these estimates is very low because of the lack of data from crustal rock-hosted systems, which are the most voluminous component of the subsurface biosphere (Colwell *et al*., 2013). Because it is predicted that rock-hosted systems have not only the space but also the energy to potentially support the metabolic activity of a large amount of biomass (Santelli *et al*., 2008), the study of these systems may provide an intriguing glimpse into unstudied forms of life and into biospheres that may exist independently of the sun.

Subsurface microbial communities associated with the geochemical process of serpentinization are of particular interest for the investigation of photosynthesis-independent ecosystems because serpentinization releases copious quantities of microbial fuel in the forms of hydrogen gas and organic carbon (Schrenk *et al*., 2013).

Serpentinization occurs at sites where tectonic action has exposed ultramafic rock from the Earth’s mantle, and it is driven by the reaction of the mineral olivine with liquid water (Schrenk *et al*., 2013). This reaction is exothermic and releases large amounts of H_2_, which is an excellent source of chemical energy for microorganisms (Proskurowski, 2010). In the presence of carbonate or dissolved CO_2_, serpentinization provides the appropriate conditions for abiogenic hydrocarbon production (Proskurowski, 2010). Abiogenic organic carbon found at sites of active serpentinization is primarily in the form of methane, but other small hydrocarbons and organic acids are also present (Morrill *et al*., 2014; Proskurowski, 2010; Lang *et al*., 2010). Effectively, serpentinization provides the two most basic requirements for life: energy and biologically available carbon (Brazelton *et al*., 2013, Morrill *et al*., 2014, Schrenk *et al*., 2013).

Despite the attractive biological potential of serpentinite systems, this process also creates an ultra-basic (pH 11-12) environment with limited availability of nutrients and electron acceptors (Schrenk *et al*., 2013). Together, these harsh conditions may restrict the number and diversity of microorganisms capable of inhabiting these systems without nutrient input from surface sources (Schrenk *et al*., 2013).

Serpentinization occurs globally where peridotite (iron-rich, ultramafic rock from the Earth’s mantle) interacts with water. Continental sites of serpentinization occur where these rocks have been obducted to form ophiolites, which can provide sampling access to serpentinite fluids (Brazelton *et al*., 2013, Morrill *et al*., 2014). At Tablelands Ophiolite in Gros Morne National Park, Newfoundland, Canada, subsurface serpentinization creates serpentinite springs that form surface pools. These pools provide access to the subsurface fluids without the need for invasive sampling techniques such as drilling (Brazelton *et al*., 2013). These pools may serve as a convenient access point for sampling the subsurface microbial communities.

However, the identification of subsurface microorganisms in these pools is difficult for two reasons. Firstly, the surface pools are fed not only by the serpentinite spring but also by surface runoff. This runoff introduces contaminant microorganisms from surrounding surface communities. Secondly, as the hydrogen-rich subsurface fluids rise to the surface, they mix with oxygenated surface fluids, creating a mixing zone that may host its own unique microbial community (Brazelton *et al*., 2013). Because samples taken from these pools are likely to contain members of all three types of communities, distinguishing between microorganisms from the subsurface, surface, and mixing zones is a challenge.

In this study we employed a new sampling technique involving the repeated emptying and refilling of the spring-fed pool in order to more directly sample the subsurface fluids and better identify those microorganisms most likely to be members of the subsurface community. Sequencing of 16S rRNA gene amplicons and metagenomes indicated that the freshly refilled pool contains microbial communities that are significantly different from those in surface environments. These results demonstrate a promising field sampling technique for the investigation of spring-fed pools and are consistent with the existence of a distinct subsurface microbial ecosystem at the Tablelands.

## Materials and Methods

### Site Description

Fluid and sediment samples were collected from the Tablelands Ophiolite ultrabasic serpentinite spring WCH2 located in Gros Morne National Park, Newfoundland Canada (N 49°27’58.7” W 057°57’29.2”). WHC2 is a shallow surface pool approximately 130 cm wide and 40 cm deep (Morrill et al., 2014). The pool is fed both by a serpentinite spring and by surface runoff. Previous work identified three sample sites (A, B, and C) within the pool (Brazelton *et al*., 2013). This project only used samples collected from site B.

### Sample Collection

Sediment samples were collected via sterile syringe from a location in the pool where previous work has identified the most likely source of subsurface fluids. Sediment was allowed to settle, and sediment-free water was expelled from the syringe. Water samples were collected for DNA sequencing on 0.2 μm Sterivex (Millipore) filter cartridges connected in-line with tubing (flushed with the sample prior to filtration) through which water was pumped with a portable peristaltic pump. Sterivex filter cartridges were placed on ice in the field and frozen in liquid nitrogen within a few hours.

Prior to any disruption, the pool was sampled by placing the intake of the tubing at the bottom of the pool near the inferred source of subsurface fluids. The pool was then emptied using the peristaltic pump, and sources of surface runoff into the pool were physically blocked and diverted with a siphon. The pool was allowed to refill for 45 minutes with water from the spring before being emptied once again. This empty-and-refill procedure was carried out three times, and then water samples were collected from the source of the subsurface fluids as the pool refilled. The process of emptying the pool and refilling three times followed by sampling was carried out three times, once each day for three days. Dissolved oxygen (DO), oxidation-reduction potential (ORP), pH, and temperature measurements were taken using a YSI Professional Plus multiparameter meter connected in-line with the peristaltic tubing, and readings were collected prior to each sample collection.

### DNA Extraction

#### Water Samples

DNA extractions were performed according to a previously published protocol (Huber *et al*. 2002; Sogin *et al*., 2006) with a few modifications described here. Chemical lysis was carried out using lysozyme, mutanolysin, proteinase K, and SDS. Bead beating was used for physical lysis. Phenol/chloroform clean-up was followed by ethanol precipitation with the addition of glycogen to increase DNA yield. DNA was resuspended in low EDTA-TE (10 mM Tris-HCl and 0.1 mM EDTA) and stored at −20°C.

#### Sediment Samples

DNA from sediment samples was extracted using PowerSoil DNA Isolation Kit (MO BIO Laboratories) as per manufacturer’s instructions and stored at −20°C.

#### DNA Cleanup

DNA samples were washed with 3 mL 65°C 10 mM Tris-HCl, pH 8, in Amicon 50K columns. Samples that did not amplify in downstream PCR reactions were further cleaned using the Boreal Genomics Aurora DNA purification system.

#### Bacterial 16S rRNA gene sequencing and processing

Bacterial 16S rRNA gene amplicon sequencing was conducted by the Michigan State University genomics core facility. The V4 region of the 16S rRNA gene (defined by primers 515F/806R) was amplified with dual-indexed Illumina fusion primers as described by Kozich *et al*., (2010). Amplicon concentrations were normalized and pooled using an Invitrogen SequalPrep DNA Normalization Plate. After library QC and quantitation, the pool was loaded on an Illumina MiSeq v2 flow cell and sequenced using a standard 500 cycle reagent kit. Base calling was performed by Illumina Real Time Analysis (RTA) software v1.18.54. Output of RTA was demultiplexed and converted to FastQ files using Illumina Bcl2fastq v1.8.4.

Paired-end sequences were filtered and merged with USEARCH 8 (Edgar, 2010), and additional quality filtering was conducted with the mothur software platform (Schloss *et al*., 2009) to remove any sequences with ambiguous bases and more than 8 homopolymers. Chimeras were removed with mothur’s implementation of UCHIME (Edgar *et al*., 2011). The sequences were pre-clustered with the mothur command pre.cluster (diffs=1), which reduced the number of unique sequences from 513,630 to 409,663. These unique sequences formed the basis of all alpha and beta diversity analyses and, therefore, were considered to be “species” for the purposes of this study. We chose not to cluster sequences any more broadly into operational taxonomic units because clustering inevitably results in a loss of biological information and because no arbitrary sequence similarity threshold can be demonstrated to consistently correspond to species-like units.

Taxonomic classification of all sequences was performed with mothur using the SILVA reference alignment (SSURefv123) and taxonomy outline (Pruesse *et al*., 2012). Differences in the relative abundances of sequences between groups of samples were measured with the aid of the R package edgeR (Robinson et al., 2010) as recommended by McMurdie *et al*., (2014). Taxonomic counts generated by mothur and edgeR results were visualized in bar charts generated with the aid of the R package phyloseq (McMurdie *et al*., 2013). Multi-dimensional scaling (MDS) plots were generated from a table of unique sequence abundances across samples (count_table from mothur) using the distance, ordinate, and plot_ordination commands in phyloseq. Raw counts were converted to proportions to normalize for variations in sequence numbers among samples prior to beta diversity calculations. Dissimilarity was partitioned into nestedness and species turnover with the R package betapart (Baselga *et al*., 2012).

### Metagenome Library Preparation and Sequencing

50 ng of DNA was suspended in a volume of 55.5 μL and sonicated at 4°C, 25% amplitude, with a 10 second pulse for 1 minute in a Qsonica Q800R2 sonicator. Each sample was divided by magnetic ‘speed bead’ size selection into large (<800bps) and small (500-800bps) subsets (for preparation of speed beads, see Rohland *et al*., 2012). For right-side selection, 0.55x of speed beads were added to 55.5 μL of sheared sample. Large fragments, bound to beads, were eluted in 55.5 μL of low EDTA-TE. Small fragments retained in supernatant were rebound using 1.8x speed beads and eluted in 55.5 μL of low EDTA-TE.

Small and large fragment libraries were prepared for Illumina sequencing using the NEBNext Q5 Hot Start HiFi PCR Master Mix with the following modifications to kit instructions: Part 1.3B, ‘no size selection’, was used. Instead, 86.5μL speed beads were used for the post-PCR clean-up. Libraries were stored at −20°C prior to sequencing with a MiSeq platform (250 Cycle Paired-End sequencing v2) at the University of Utah High-Throughput Genomics core facility. Data generated by the platform was processed through the CASAVA version 1.8 pipeline.

Environmental shotgun DNA sequences for each sample were preprocessed and assembled separately using an in-house pipeline. Artifactual sequences were identified and removed with cutadapt (Martin, 2011). Reads found to have Illumina adapters starting at their 5’-end were discarded, and reads containing Illumina adapters towards the 3’-end were trimmed where the first adapter began. Identical and 5’-prefix replicates were removed as suggested by Gomez-Alvarez *et al*. (2009). For some datasets, positions at the start or end of the read would exhibit nucleotide frequencies inconsistent with the rest of the sequence content. When this occurred, reads were cropped at the discrepant position. Low-quality bases were removed from the ends of the reads using an in-house script. The remaining sequence was scanned 20 base pairs at a time and trimmed where the mean quality score fell below the threshold. A threshold of 8 was used for all quality-based trimming steps. Reads that did not pass a minimum length threshold of 62 base pairs after quality and adapter trimming were removed from the dataset. All in-house scripts used in the pipeline are freely available and can be found at https://www.github.com/Brazelton-Lab/seq_qc.

Large and small insert libraries were preprocessed individually and then co-assembled with Ray Meta (Boisvert, 2012). High-quality reads were mapped onto the selected assembly with Bowtie2 (Langmead, 2012), and insert-coverage for mis-assembly detection was determined using bedtools genomecov (Quinan, 2010) with the flag -pc to indicate that coverage should be determined by insert instead of read. Mis-assemblies were detected as in Sharon et al (2013). The Prokka pipeline (Seeman, 2014) was used for gene prediction and functional annotation. The arguments --metagenome and --proteins were used with Prokka to indicate that genes should be predicted with the implementation of Prodigal (Hyatt, 2010) optimized for metagenomes, and then searched preferentially against a user-provided database. The protein database provided was the last free version of the Kyoto Encyclopedia of Genes and Genomes (Ogata, 1999). Metabolic pathways present in the community were determined by providing a table containing normalized counts of orthologous genes to HUMAnN2 (Abubucker, 2012). The pathway mapping used with HUMAnN2 was obtained from the FOAM ontology (Prestat, 2014). Percent coverage of metabolic pathways was calculated as the proportion of reads per kilobase (RPK) for each metabolic pathway to total RPK.

## Results

### 16S rRNA Genetic Diversity

We collected twenty-five DNA samples from the field. These included replicates from the undisturbed pool, sediments from the bottom of the pool, surface fluids samples, and samples from each of the three refills (Table 1). High coverage Illumina MiSeq sequencing of bacterial 16S rRNA gene amplicons resulted in a total of 3,064,527 sequences. These sequences were used to compare community dissimilarity among samples (calculated from abundance distributions of unique sequences) and for taxonomic classification of bacterial communities.

**Table 1:**
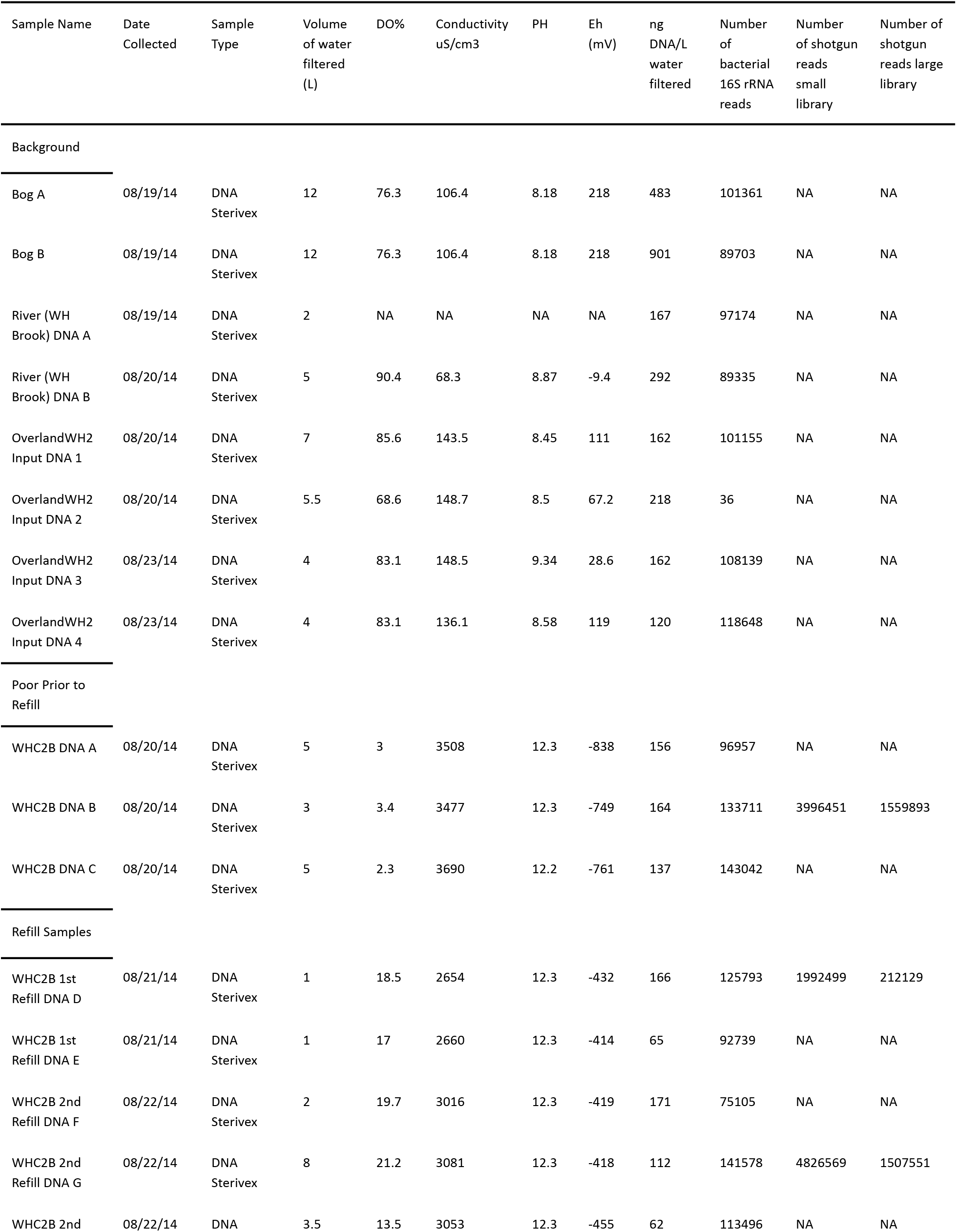
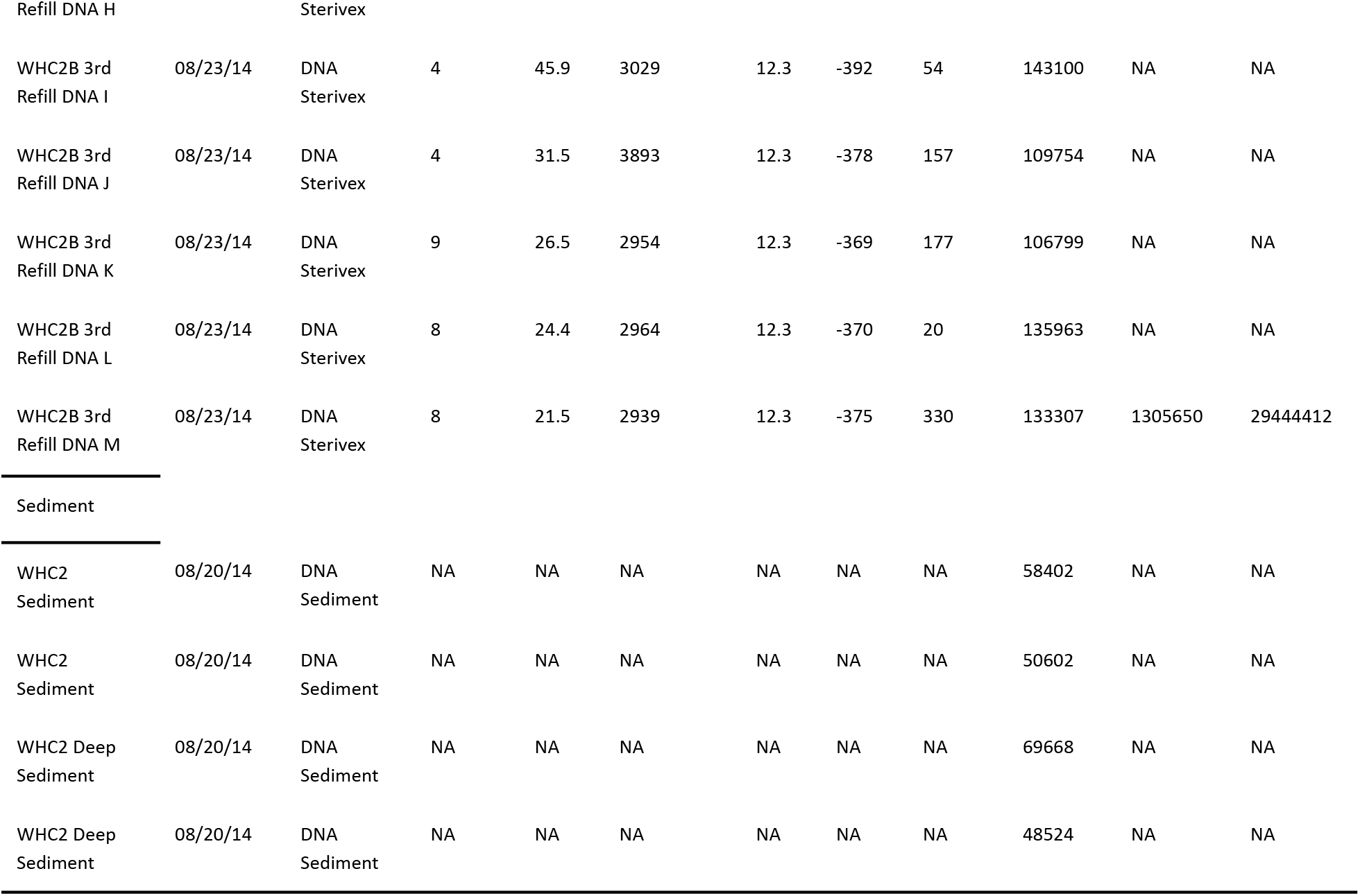
Sample Details.

| Sample Name | Date Collected | Sample Type | Volume of water filtered (L) | DO% | Conductivity uS/cm3 | PH | Eh (mV) | ng DNA/L water filtered | Number of bacterial 16S rRNA reads | Number of shotgun reads small library | Number of shotgun reads large library |
| --- | --- | --- | --- | --- | --- | --- | --- | --- | --- | --- | --- |
| Background |  |  |  |  |  |  |  |  |  |  |  |
| Bog A | 08/19/14 | DNA Sterivex | 12 | 76.3 | 106.4 | 8.18 | 218 | 483 | 101361 | NA | NA |
| Bog B | 08/19/14 | DNA Sterivex | 12 | 76.3 | 106.4 | 8.18 | 218 | 901 | 89703 | NA | NA |
| River (WH Brook) DNA A | 08/19/14 | DNA Sterivex | 2 | NA | NA | NA | NA | 167 | 97174 | NA | NA |
| River (WH Brook) DNA B | 08/20/14 | DNA Sterivex | 5 | 90.4 | 68.3 | 8.87 | -9.4 | 292 | 89335 | NA | NA |
| OverlandWH2 Input DNA 1 | 08/20/14 | DNA Sterivex | 7 | 85.6 | 143.5 | 8.45 | 111 | 162 | 101155 | NA | NA |
| OverlandWH2 Input DNA 2 | 08/20/14 | DNA Sterivex | 5.5 | 68.6 | 148.7 | 8.5 | 67.2 | 218 | 36 | NA | NA |
| OverlandWH2 Input DNA 3 | 08/23/14 | DNA Sterivex | 4 | 83.1 | 148.5 | 9.34 | 28.6 | 162 | 108139 | NA | NA |
| OverlandWH2 Input DNA 4 | 08/23/14 | DNA Sterivex | 4 | 83.1 | 136.1 | 8.58 | 119 | 120 | 118648 | NA | NA |
| Poor Prior to Refill |  |  |  |  |  |  |  |  |  |  |  |
| WHC2B DNA A | 08/20/14 | DNA Sterivex | 5 | 3 | 3508 | 12.3 | -838 | 156 | 96957 | NA | NA |
| WHC2B DNA B | 08/20/14 | DNA Sterivex | 3 | 3.4 | 3477 | 12.3 | -749 | 164 | 133711 | 3996451 | 1559893 |
| WHC2B DNA C | 08/20/14 | DNA Sterivex | 5 | 2.3 | 3690 | 12.2 | -761 | 137 | 143042 | NA | NA |
| Refill Samples |  |  |  |  |  |  |  |  |  |  |  |
| WHC2B 1st Refill DNA D | 08/21/14 | DNA Sterivex | 1 | 18.5 | 2654 | 12.3 | -432 | 166 | 125793 | 1992499 | 212129 |
| WHC2B 1st Refill DNA E | 08/21/14 | DNA Sterivex | 1 | 17 | 2660 | 12.3 | -414 | 65 | 92739 | NA | NA |
| WHC2B 2nd Refill DNA F | 08/22/14 | DNA Sterivex | 2 | 19.7 | 3016 | 12.3 | -419 | 171 | 75105 | NA | NA |
| WHC2B 2nd Refill DNA G | 08/22/14 | DNA Sterivex | 8 | 21.2 | 3081 | 12.3 | -418 | 112 | 141578 | 4826569 | 1507551 |
| WHC2B 2nd | 08/22/14 | DNA | 3.5 | 13.5 | 3053 | 12.3 | -455 | 62 | 113496 | NA | NA |

Dissimilarity among whole bacterial community compositions (as measured by the Bray-Curtis index calculated from the abundance distributions of unique sequences among all samples) was visualized with a multi-dimensional scaling (MDS) plot (Figure 1). Bacterial communities from the refills clustered together (95% confidence interval) in the MDS plot, reflecting that their sequence compositions are significantly more similar to each other than to any other samples. All refills were more similar to the pool prior to disruption than to any other samples, and the MDS plot highlights strong contrasts among the bacterial community compositions of the pool samples compared to sediment and surface water samples (Figure 1).

**Figure 1:**
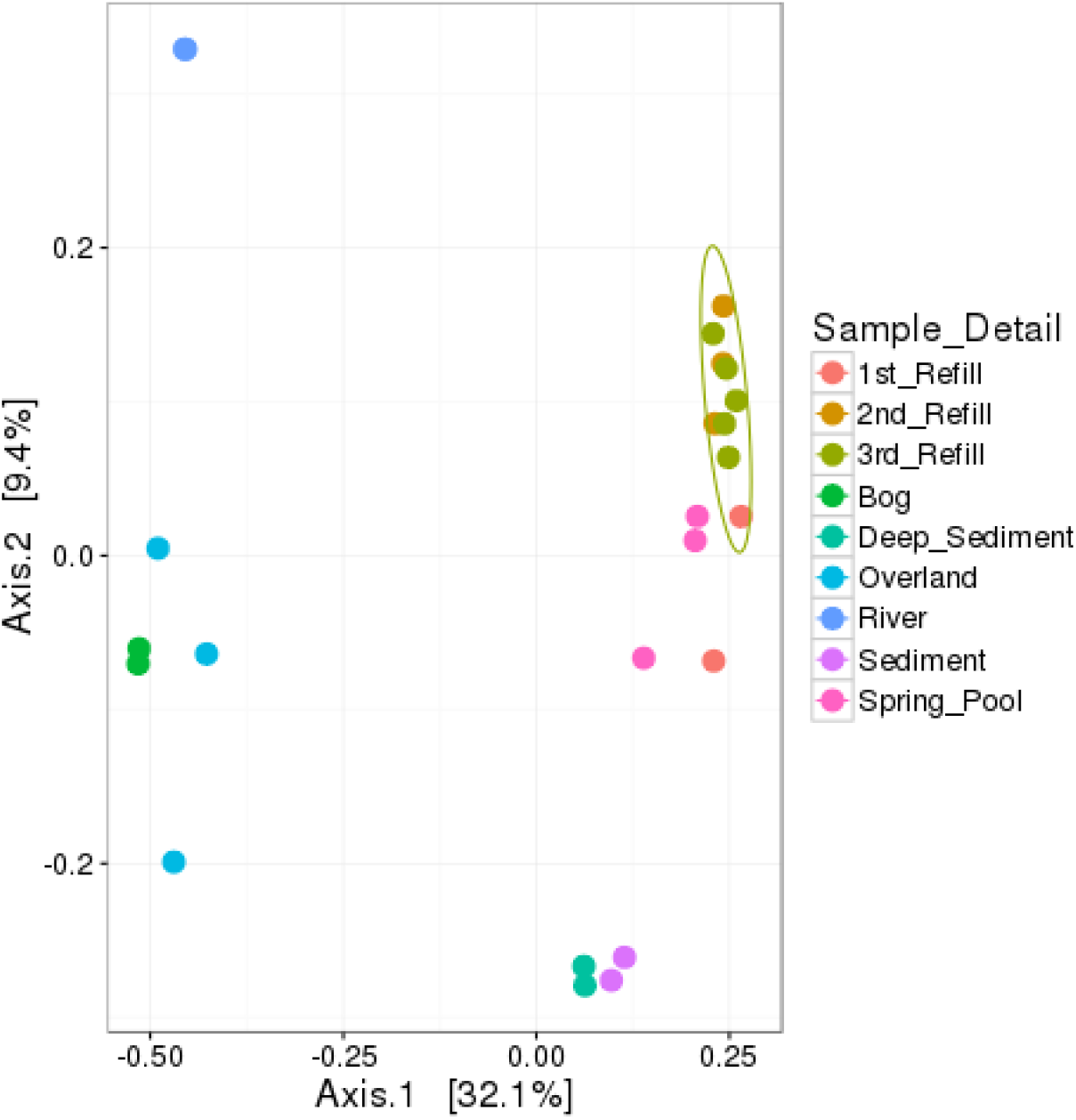
Bacterial community composition dissimilarity (as calculated with the Bray-Curtis index) as measured by environmental 16S rRNA gene amplicon sequencing. Each point represents the whole bacterial community composition of one environmental sample.

Community dissimilarity values were partitioned into nestedness and turnover components in order to test whether the observed differences in beta diversity were due to loss/gain of species (nestedness) or to replacing species with other species (turnover) (Baselga *et al*., 2012). (‘Species’ are defined as unique, quality-filtered, and pre-clustered sequences for the purposes of this study, as explained in the methods). The average of these dissimilarity values and their nestedness and turnover components are reported in Table 2. For example, of the 0.84-0.87 Sorenson dissimilarity between bacterial community compositions of the pool prior to disruption and those of the third refill, 85-99% of this dissimilarity was due to turnover, while nestedness accounted for only 0.1-15% of the dissimilarity.

**Table 2:** Average Dissimilarity Values.

| Sample Comparison | Average Bray<br>Curtis Dissimilarity | Average Sorenson<br>Dissimilarity | Average Simpson<br>Dissimilarity<br>(turnover) | Average<br>Dissimilarity due<br>to Nestedness |
| --- | --- | --- | --- | --- |
| Pool to Third Refill | 0.56 | 0.85 | 0.81 | 0.04 |
| Sediment to Third Refill | 0.64 | 0.91 | 0.88 | 0.03 |
| First Refill to Third Refill | 0.46 | 0.82 | 0.79 | 0.03 |
| Second Refill to Third Refill | 0.43 | 0.82 | 0.79 | 0.03 |

### Taxonomic Classification of Bacterial Communities

Differences in bacterial community composition between the pool prior to disruption and the third refill were examined in more detail by conducting a differential abundance analysis with these samples. The pool was represented by 3 field replicates, and the third refill was represented by 5 field replicates (Table 1). In Figure 2, each data point of the plot represents the difference in abundance of a single sequence between the pool prior to disruption and the third refill. The plot’s x-axis displays the mean abundance (in units of log2 counts per million) of each sequence across all 8 samples (3 pool replicates and 5 third refill replicates). Red data points indicate sequences whose differential abundance between the two sample types passed a significance test (p-value < 0.05 and false discovery rate < 0.05) implemented by the edgeR package (Robinson *et al*., 2010).

**Figure 2:**
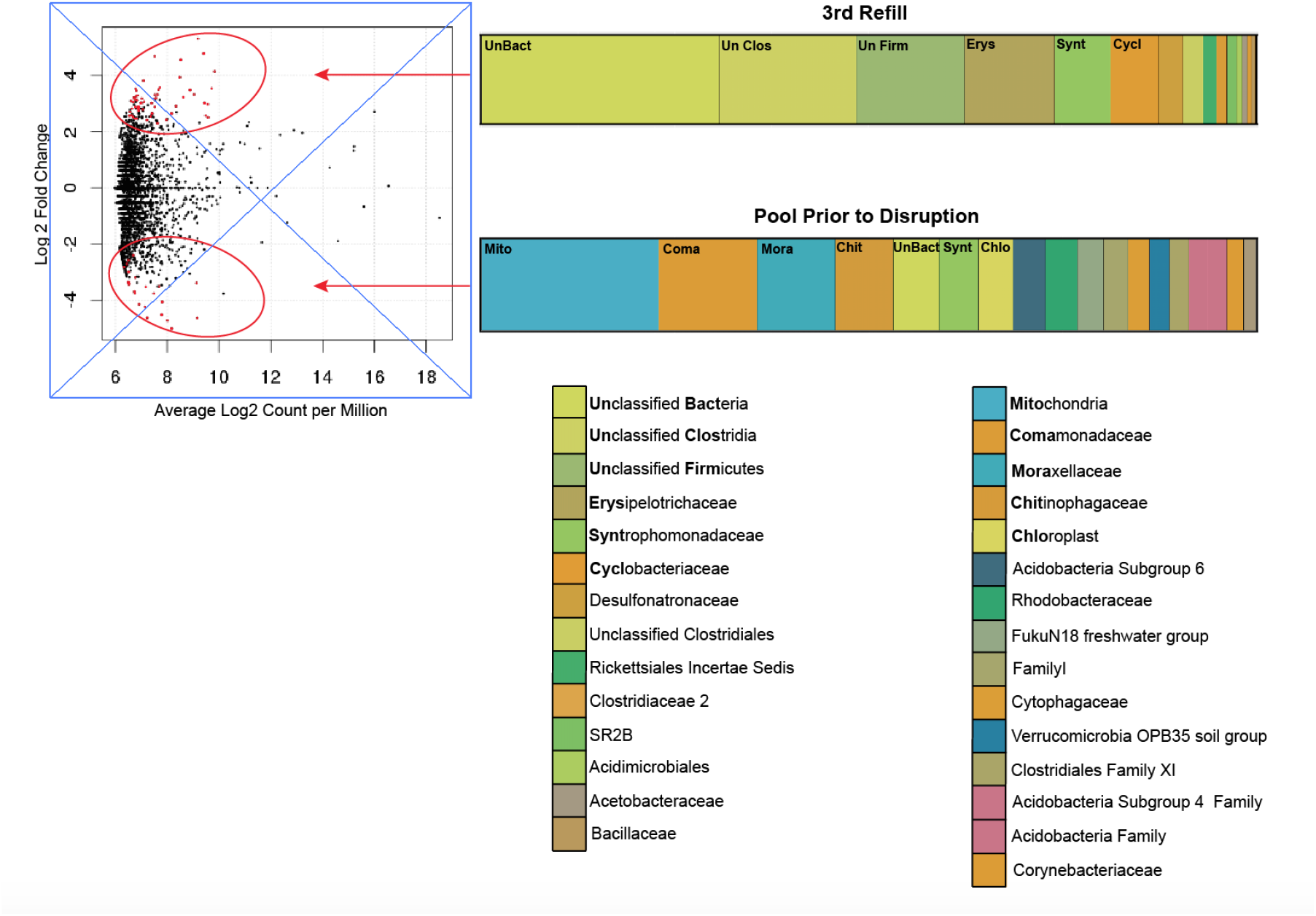
Taxonomic classifications of bacterial 16S rRNA gene sequences that were identified as significantly enriched in the third refill compared to the pool prior to disruption, and vice versa. Red dots indicate significantly enriched sequences with less than 0.05 false discovery rates. Bar charts group the enriched sequences according to their taxonomic classifications at a family level.

Many sequences representing mitochondria and chloroplasts, expected to represent surface contamination by Eukarya, were significantly enriched in the samples from the pool prior to disruption. Compatible with previous experiments at this location, *Erysipelotrichaceae* was enriched in the refill samples (Brazelton *et al*., 2013). The vast majority of refill-enriched sequences could only be classified at the domain, phylum, or class level, suggesting that these are novel taxa or at least poorly represented in taxonomic databases. Both the pool-enriched and refill-enriched subsets include sequences that were classified as *Syntrophomonadaceae*. Note that the sequences classified as *Syntrophomonadaceae* enriched in the pool are distinct from the sequences classified as *Syntrophomonadaceae* enriched in the refill, as it is not possible for the same sequence to be enriched in both sample types.

### Metagenomic Analysis

To explore the differences in metabolic potential between the refills and the undisturbed pool, metagenomic analyses were carried out on one sample from the undisturbed pool (DNA B) and one sample from each of the three refills (DNA D, DNA G, DNA M) (see Table 1 for sample details). For these analyses, shotgun metagenomic sequences from the selected samples were assembled into contiguous genomic fragments (contigs), which were used for gene prediction, annotation, and categorization of metabolic pathways. Predicted proteins that were able to be assigned a KEGG ID were grouped into metabolic pathways according to the FOAM ontology of pathways (Prestat 2014).

Table 3 lists the fifteen most abundant pathways in the third refill and the fifteen most abundant pathways in the pool prior to disruption (as measured by percent coverage). Homoacetogenesis and the reductive TCA cycle featured more prominently in the refills than in the undisrupted pool. Both of these pathways are consistent with non-photosynthetic carbon fixation in hydrogen rich, anaerobic environments (Diekert *et al*., 1994; Ragsdale *et al*., 2008). The aerobic hydrogen oxidation pathway enriched in the pool prior to disruption is consistent with the presence of microorganisms that capitalize on the mix of hydrogen-rich subsurface fluids and oxygen-rich surface fluids present in the undisturbed pool.

**Table 3:**
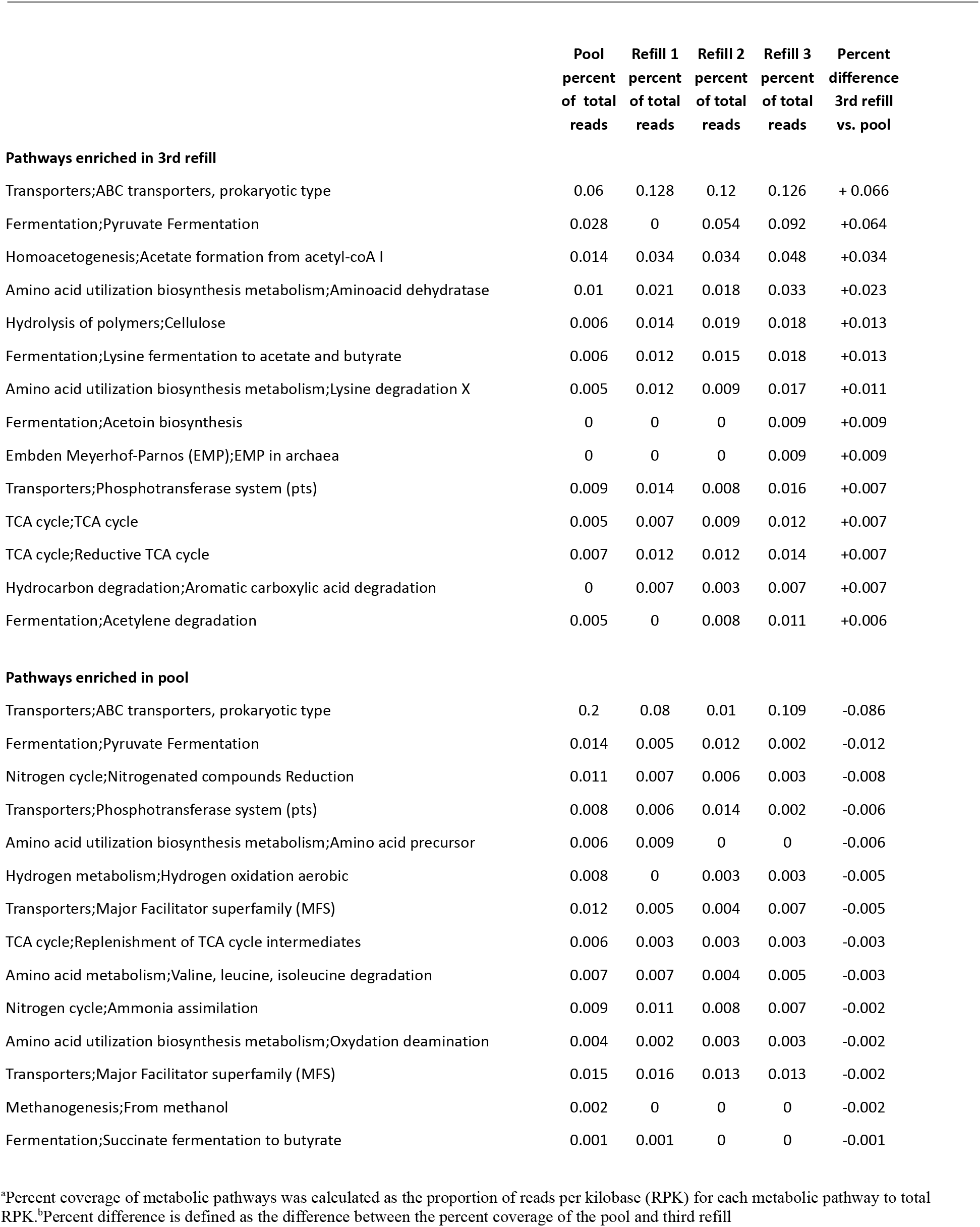
Metabolic Pathways.

The ABC transporters and pyruvate oxidation pathways listed as enriched in both sets of samples, are distinct pathways at finer resolution (i.e. different forms of pyruvate oxidation), but have been grouped for table simplicity.

## Discussion

One of the primary objectives of this project was to create and validate a sampling procedure that allowed for more direct sampling of the subsurface fluids. The process of emptying and refilling the pool only from the subsurface source appears to have accomplished this goal. The microbial composition of the refill samples was significantly different from the microbial composition of surface sources, suggesting that the freshly refilled pool is more representative of subsurface habitats (Figure 1). This conclusion is further supported by the lack of nestedness contributing to the dissimilarity between the communities in the refill samples and other sample types. Had the refill samples been subsets of the typical pool community or of the sediment found at the bottom of the pool, we would expect the nestedness component of beta diversity to be the primary contributor to overall community dissimilarity. In that case, the refill samples could be seen as smaller subsets of the sediments or surface-based communities rather than representative of a distinct subsurface community. However, the opposite trend is observed, thus indicating that the samples taken from the refills represent a distinct community.

These results highlight specific microbial taxa that may be key members of a subsurface, serpentinization-associated ecosystem. For example, *Erysipelotrichaceae* were shown to be well-represented in the refill samples, consistent with our previous work that identified *Erysipelotrichaceae* as the most likely representatives of a subsurface, serpentinization-associated ecosystem (Brazelton *et al*., 2013).

The vast majority of sequences enriched in the refill samples could not be classified below a phylum or class level, so our best insights into the nature of these potential subsurface inhabitants come from predictions of metabolic pathways from shotgun metagenomic data. The most abundant carbon fixation pathways (homoacetogenesis and the reverse TCA cycle) suggest that the microbial inhabitants of the subsurface ecosystem underlying the Tablelands serpentinite springs are able to fix carbon from dissolved inorganic carbon using hydrogen as an electron donor - these substrates exist in excess in serpentinite-hosted subsurface systems (Proskurowski 2010). Fermentation pathways and the TCA cycle (which can function anaerobically in bacteria (Alteri *et al*., 2012)) are enriched in the refill samples and are consistent with the metabolism of small hydrocarbons.

The same work that identified *Erysipelotrichaceae* as a likely member of the subsurface also identified the betaproteobacterial genus *Hydrogenophaga* as an inhabitant of the mixing zone (Brazelton *et al*., 2013). These organisms are thought to use hydrogen oxidation to fix carbon when organic carbon is not present (Willems *et al*., 1989). However, because they are aerobic or facultatively anaerobic it is likely that they live in the transition zone where both hydrogen-rich subsurface fluids and oxygenated surface fluids are present but unlikely that they would be found in anoxic samples closely associated with the subsurface (Brazelton *et al*., 2013; Willems *et al*., 1989). Our metagenomic results, showing aerobic hydrogen oxidation present in the pool prior to disruption but diminished in the refills (Table 2), are consistent with this interpretation.

Methanogenesis pathways were notably absent from the metagenomic results reported here, consistent with previous work at the Tablelands (Brazelton *et al*., 2013) and in strong contrast to the prevalence of methanogens in deep-sea chimneys of the Lost City hydrothermal field (Schrenk *et al*., 2013). At this site, carbonate chimneys host thick biofilms heavily populated by a single phylotype of Methanosarcinales (Schrenk *et al*., 2004). The apparent absence of methanogens at the Tablelands is a potentially important clue to understanding key differences between continental and marine sites of serpentinization.

The distinctive metabolic potential of organisms inferred to have subsurface origins is consistent with the possibility that these organisms live independently of photosynthesis. However, DNA-based analyses cannot indicate metabolic activity in situ, and the subsurface-associated microbes identified here may represent dead or dormant microorganisms. To address this question, future studies will conduct RNA sequencing and metabolic activity experiments. If the communities shown to be most closely associated with the subsurface are also shown to be metabolically active, it will further support the idea that serpentinite-hosted systems are capable of supporting a thriving biosphere independent of the sun.

The results from our study show that by emptying and refilling the pool multiple times prior to sampling, we were better able to identify subsurface microbes. Employing a similar technique in future studies at other sites of continental serpentinization should help to better characterize the benefits and challenges presented to life by serpentinization.

